# Cultivar-specific transcriptome and pan-transcriptome reconstruction of tetraploid potato

**DOI:** 10.1101/845818

**Authors:** Marko Petek, Maja Zagorščak, Živa Ramšak, Sheri Sanders, Špela Tomaž, Elizabeth Tseng, Mohamed Zouine, Anna Coll, Kristina Gruden

## Abstract

Although the reference genome of *Solanum tuberosum* Group Phureja double-monoploid (DM) clone is available, knowledge on the genetic diversity of the highly heterozygous tetraploid Group Tuberosum, representing most cultivated varieties, remains largely unexplored. This lack of knowledge hinders further progress in potato research. In conducted investigation, we first merged and manually curated the two existing partially-overlapping DM genome-based gene models, creating a union of genes in Phureja scaffold. Next, we compiled available and newly generated RNA-Seq datasets (cca. 1.5 billion reads) for three tetraploid potato genotypes (cultivar Désirée, cultivar Rywal, and breeding clone PW363) with diverse breeding pedigrees. Short-read transcriptomes were assembled using several *de novo* assemblers under different settings to test for optimal outcome. For cultivar Rywal, PacBio Iso-Seq full-length transcriptome sequencing was also performed. EvidentialGene redundancy-reducing pipeline complemented with in-house developed scripts was employed to produce accurate and complete cultivar-specific transcriptomes, as well as to attain the pan-transcriptome. The generated transcriptomes and pan-transcriptome represent a valuable resource for potato gene variability exploration, high-throughput omics analyses, and breeding programmes.

## Background & Summary

At species level, genomes of individuals can differ in single nucleotide polymorphisms (SNPs), short insertions and deletions (INDELs), gene copy numbers, and presence or absence of genes [1]. The latter leads to the concept of species specific pan-genomes, namely the core genome present in most individuals and the dispensable genome comprised of genes present only in a subset of individuals, which results in the emergence of particular subgroup-specific phenotypes. This concept has been extended to pan-transcriptomes, where the presence or absence of variations is not bound only to gene content, but also to genetic and epigenetic regulatory elements. Pan-genomes and pan-transcriptomes have been described in the model plant species *Arabidopsis thaliana* [2] and several crop species, including maize [3, 4], rice [5], wheat [6] and soybean [7].

While the genome of a double-monoploid clone of *Solanum tuberosum* Group Phureja (DM) is available [8], this diploid potato group differs from the tetraploid Group Tuberosum, which includes most varieties of cultivated potato. Through domestication and modern breeding efforts, different potato cultivars also acquired genes from other closely related Solanum species or lost some ancestral genes [1]. Different breeding programmes have resulted in accumulation of smaller genome modifications, e.g. SNPs and INDELs. Consequently, each distinct potato cultivar harbours a unique set of transcripts, resulting in physiological and developmental differences and different responses to biotic and abiotic stress. SNP and INDEL profile differences and novel gene variants in anthocyanin pathway were identified in a comparative transcriptome analysis of two Chinese potato cultivars [9]. Unfortunately, we could not include these transcriptomes in our pan-transcriptome because the assemblies were not publicly accessible.

Based on the DM genome, the PGSC and ITAG annotation consortia [8, 10] have each independently produced potato gene models. For practical reasons, most potato researchers use only one genome annotation, either PGSC or ITAG, especially when conducting high-throughput analyses. Using an incomplete gene set can lead to false conclusions regarding gene presence or gene family diversity in potato. Using a computational pipeline followed by manual curation, we have consolidated the two publicly available Group Phureja DM gene model sets to produce a unified one.

While a combined DM gene set is useful, it is still not as useful as a pan-transcriptome that includes assemblies from cultivated potatoes. However, obtaining an optimal transcriptome from short-read RNA-Seq data is not a trivial task. Each *de novo* assembler suffers from different intrinsic error generation and no single assembler performs best on all datasets [11]. To maximise diversity and completeness of potato cultivar transcriptomes, usage of multiple *de novo* transcriptome assemblers and various parameter combinations over the same input data was employed. Following this “overassembly” step, we used tr2aacds pipeline from EvidentialGene [12] to reduce redundancy across assemblies and obtain cultivar-specific transcriptomes. Finally, we consolidated representative cultivar-specific sequences to generate a potato pan-transcriptome (StPanTr). These transcriptomes will improve high throughput sequencing analyses, from RNA-Seq and sRNA-Seq to more specific ones like ATAC-Seq, by providing a more comprehensive and accurate mapping reference. The translated protein sequences can enhance the sensitivity of high-throughput mass-spectroscopy based proteomics. The resource is valuable also for the design of various PCR assays, e.g. quantitative PCR, where exact sequence information is required. Additionally, the knowledge generated regarding variations in transcript sequences between cultivars, such as SNPs, insertions and deletions, will be a key instrument in assisting potato breeding programmes.

### Data records

Transcriptomic sequences of three potato genotypes, cv. Désirée, cv. Rywal and breeding clone PW363, were obtained from in-house RNA-Seq projects [13–17] and supplemented by publicly available cv. Désirée datasets retrieved from SRA [18–20] (Table 1). All generated files have been deposited to FAIRDOMhub [21] under project name _p_stRT (https://fairdomhub.org/projects/161).

**Table 1.** Samples used to generate the *de novo* transcriptome assemblies.

| Genotype | Sample description<sup>1</sup> | Sequencing platform | Library type<sup>2</sup> | Number of reads<sup>3</sup> | SRA ID |
| --- | --- | --- | --- | --- | --- |
| Désirée | PVY inoculated leaves | Illumina | DSN-normalized PE90 unstranded | ~54 mio | SRR10070125 |
| Désirée | non-transformed and PVY-inoculated plants, non-infested and CPB infested leaves | Illumina | PE90 unstranded | ~195 mio | SRR1207287, ..., SRR1207290 |
| Désirée | mock and PVY inoculated leaves and stem | SOLiD | SE50 unstranded | ~154 mio | SRR10065428, SRR10065429 |
| Désirée | leaves | Illumina | SE50 unstranded | ~172 mio | SRR3161991, SRR3161995, SRR3161999, SRR3162003, SRR3162007, SRR3162011, SRR3162015, SRR3162019, SRR3162023, SRR3162027, SRR3162031, SRR3162035 |
| Désirée | seedlings | Illumina | SE100 unstranded | ~80 mio | SRR4125238, ..., SRR4125247 |
| Désirée | roots | Illumina | SE100 unstranded | ~31 mio | SRR4125248, ..., SRR4125252 |
| Désirée | mock and *Phytophthora infestans* inoculated leaves | Illumina | PE90 unstranded | ~53 mio | ERR305632 |
| Rywal | mock and PVY inoculated leaves | PacBio | Iso-Seq, 0.7–2 Kb, 2–3.5 Kb, >3.5 Kb | ~1.4 mio CCS | SRR8281993, ..., SRR8282008 |
| Rywal | mock and PVY inoculated leaves | Illumina | PE100 strand-specific | ~710 mio | SRX6801457, ..., SRX6801468 |
| PW363 | PVY inoculated leaves | Illumina | DSN-normalized PE90 unstranded | ~104 mio | SRR10070123, SRR10070124 |
| PW363 | mock and PVY inoculated leaves | SOLiD | SE50 unstranded | ~180 mio | SRR10065430, ..., SRR10065433 |

<sup>1</sup> PVY, *Potato virus Y*; CPB, Colorado potato beetle
<sup>2</sup> PE, paired-end library (the number stands for read length in nt); SE, single-end library (the number stands for read length in nt); DSN-normalized, RNA-Seq library utilizing the crab duplex nuclease; CCS, circular consensus sequences
<sup>3</sup> For paired-end libraries, pairs are counted as two reads

The largest quantity of reads, cca. 739 mio reads of various lengths, was obtained for cv. Désirée, using Illumina and SOLiD short-read sequencing platforms. For cv. Rywal and breeding clone PW363 only mature leaf samples were available. For cv. Désirée leaf samples were augmented with samples from stems, seedlings and roots. For cv. Rywal short-read sequencing was complemented with full-length PacBio Iso-Seq sequencing of independent samples. Detailed sample information is provided in Supplementary Table 1.

Cv. Rywal NPR1-1 coding sequences, sequenced by the Sanger method, were deposited at NCBI GenBank under accession numbers MT210578, …, MT210585 [22–29].

The GTF file with merged ITAG and PGSC gene models for *S. tuberosum* Group Phureja DM genome v4.04 [30] is also available at FAIRDOMHub project page, as well as the cultivar-specific and pantranscriptome assembly FASTA and annotation files [31].

## Methods

### Merging PGSC and ITAG gene models of reference genome Group Phureja

GFF files corresponding to their respective gene models (PGSC v4.04, ITAG v1.0) were retrieved from the Spud DB (solanaceae.plantbiology.msu.edu) potato genomics resource [32]. The two models (39,431 PGSC and 35,004 ITAG) were then compared on the basis of their exact chromosomal location and orientation in order to create the most complete set of genes from both predicted genome models. Several combinations arose from the merge (Figure 1a), those for which no action was required (singletons, i.e. sole PGSC or ITAG genes not covering any other genes); 1-to-1 or 1-to-2 combinations between PGSC and ITAG genes, which were solved programmatically; and lastly, combinations of more than 3 genes in various combination types, which continued on to manual curation. The latter (318 gene clusters; example in Figure 1b) were considered to be non trivial merge examples (overlapping genes in two models or multiple genes in PGSC corresponding to a single gene in ITAG). This resulted in a merged DM genome GFF3 file with 49,322 chromosome position specific sequences, of which 31,442 were assigned with ITAG gene IDs and 17,880 with PGSC gene IDs (Supplementary File 1).

**Figure 1.**
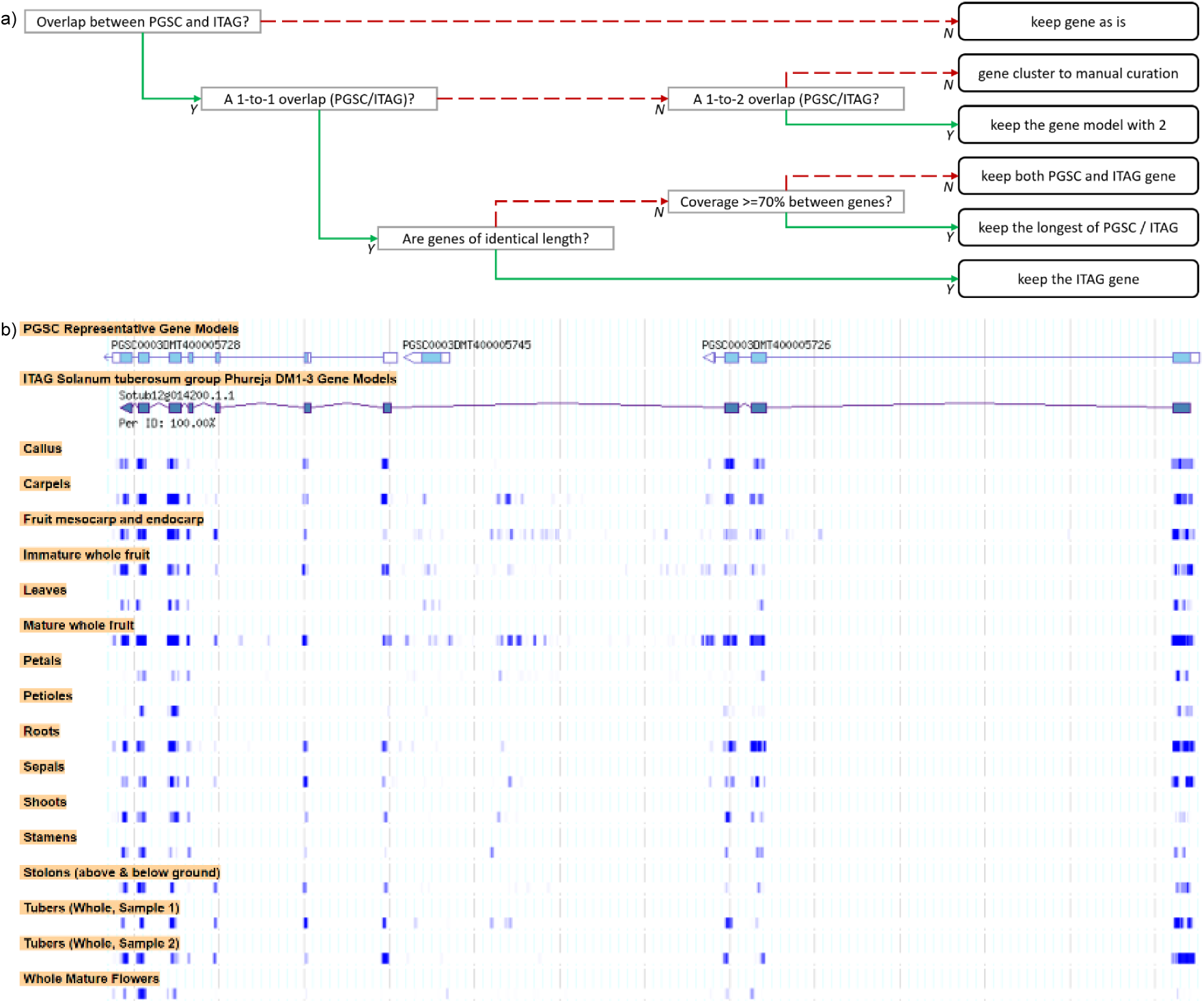
Merging DM Phureja PGSC and ITAG gene models. a) Decision tree for the merging of both genome models, with 6 possible outcomes: singleton genes (‘keep gene as is’), manual curation (‘gene cluster to manual curation’ and programmatic solution (all remaining 4 options). Green solid lines represent a satisfied condition (Y: Yes), dashed red lines, an unsatisfied condition (N: No). b) Example of manual curation in the merged DM genome GTF, region visualisation (chr12:11405699..11418575) in the Spud DB (solanaceae.plantbiology.msu.edu) Genome Browser [32]. ITAG defined Sotub12g014200.1.1 spans three PGSC defined coding sequences (PGSC0003DMT400005728, PGSC0003DMT400005745 and PGSC0003DMT400005726). Below the gene models, RNA sequence tracks are shown, showing how these genes are expressed in various plant organs. In the concrete case, Sotub12g014200.1.1 was preferred due to RNA-Seq evidence being in concordance, and no evidence for PGSC0003DMT400005745.

### Data pre-processing

The complete bioinformatic pipeline is outlined in Figure 2. Sequence quality assessment of raw RNA-Seq data, quality trimming, and removal of adapter sequences and polyA tails was performed using CLC Genomics Workbench v6.5-v10.0.1 (Qiagen) with maximum error probability threshold set to 0.01 (Phred quality score 20) and no ambiguous nucleotides allowed. Minimal trimmed sequence length allowed was set to 15bp while maximum up to 1kb. Orphaned reads were re-assigned as single-end (SE) reads. Processed reads were pooled into cultivar datasets as properly paired-end (PE) reads or SE reads per cultivar per sequencing platform. For the Velvet assembler, SOLiD reads were converted into double encoding reads using perl script “denovo_preprocessor_solid_v2.2.1.pl” [33]. To reduce the size of cv. Désirée and cv. Rywal datasets, digital normalization was performed using khmer from bbmap suite v37.68 [34] prior to conducting *de novo* assembly using Velvet and rnaSPAdes.

**Figure 2.**
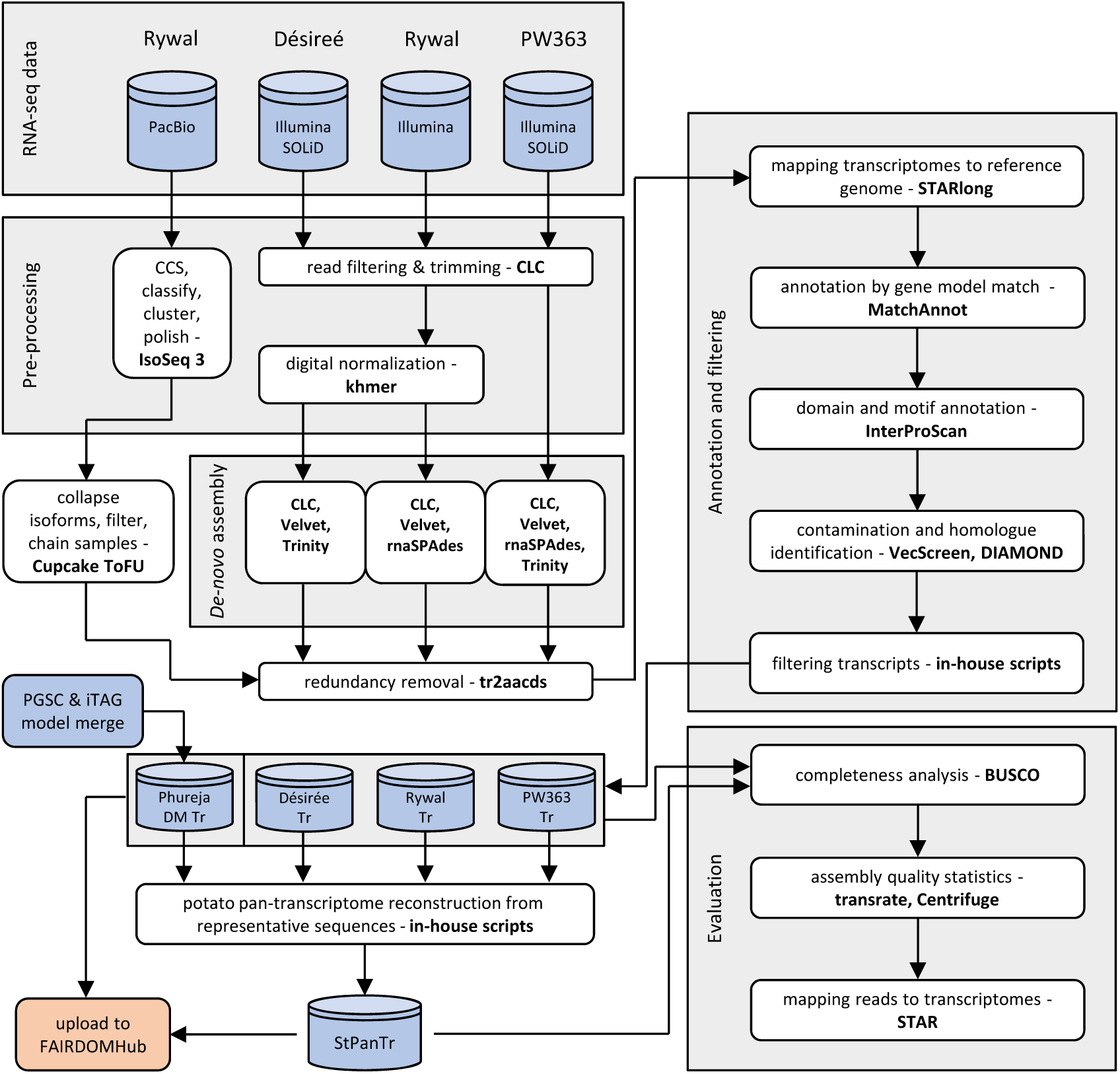
Bioinformatics pipeline for generation of potato transcriptomes. Software used in specific steps are given in bold. Input datasets (sequence reads) and output data (transcriptomes) are depicted as blue cylinders. Data upload steps to public repositories are shaded in orange. Abbreviations: SRA – NCBI Sequence Read Archive, PGSC – Potato Genome Sequencing Consortium, ITAG – international Tomato Annotation Group, CLC – CLC Genomics Workbench, PacBio – Pacific Biosciences Iso-Seq sequencing, Tr – transcriptome, StPanTr – potato pan-transcriptome, tr2aacds – “transcript to amino acid coding sequence” Perl script from EvidentialGene pipeline.

PacBio long reads were processed for each sample independently using Iso-Seq 3 analysis software (Pacific Biosciences). Briefly, the pipeline included Circular Consensus Sequence (CCS) generation, full-length reads identification (”classify” step), clustering isoforms (”cluster” step) and “polishing” step using Arrow consensus algorithm. Only high-quality full-length PacBio isoforms were used as input for further steps.

### PacBio Cupcake ToFU pipeline

Cupcake ToFU (github.com/Magdoll/cDNA_Cupcake) scripts [35] were used to further refine the Iso-Seq transcript set. Redundant PacBio isoforms were collapsed with “collapse_isoforms_by_sam.py” and counts were obtained with “get_abundance_post_collapse.py”. Isoforms with less than two supporting counts were filtered using “filter_by_count.py” and 5’-degraded isoforms were filtered using “filter_away_subset.py”. Isoforms from the two samples were combined into one non-redundant Iso-Seq transcript set using “chain_samples.py”.

### De Bruijn graph based *de novo* assembly of short reads

Short reads were *de novo* assembled using Trinity v.r2013-02-25 [36], Velvet/Oases v. 1.2.10 [37], rnaSPAdes v.3.11.1 [38] and CLC Genomics Workbench v8.5.4-v10.1.1 (Qiagen). Illumina and SOLiD reads were assembled separately. For CLC Genomics *de novo* assemblies, combinations of three bubble sizes and 14 k-mer sizes were tested on PW363 Illumina dataset. Varying bubble size length did not influence the assembly statistics much, therefore we decided to use the length of 85bp for Illumina datasets of the other two cultivars (Supplementary Figure 2). Bubble size and k-mer length parameters used for Velvet and CLC are given in Table 2. The scaffolding option in CLC and Velvet was disabled. More detailed information per assembly is provided in Supplementary Table 2.

**Table 2.** Parameters used for short read *de novo* assembly generation.

| Genotype | Assembly ID | Read type | Assembler | Assembler version | k-mer length (word size) | Bubble size |
| --- | --- | --- | --- | --- | --- | --- |
| Désirée | CLCdnDe8 | SOLiD | CLC *de novo* | 9.1 | 24 | 50 |
| Désirée | CLCdnDe1 | SOLiD | CLC *de novo* –<br>– transcript discovery<br>as reference | 10.0.1 | 24 | 50 |
| Désirée | VdnDe8, ...,<br>..., VdnDe10 | SOLiD | Velvet/Oases | 1.2.10 | 23, 33, 43 | Default |
| Désirée | CLCdnDe9, ...,<br>..., CLCdnDe14 | Illumina | CLC *de novo* | 9.1 | 21, 23, 33, 43,<br>53, 63 | 85 |
| Désirée | CLCdnDe2, ...,<br>..., CLCdnDe7 | Illumina | CLC *de novo* –<br>– transcript discovery<br>as reference | 10.0.1 | 21, 23, 33, 43,<br>53, 63 | 85 |
| Désirée | TDe | Illumina | Trinity | r2013-02-25 | 25 | NA |
| Désirée | VdnDe1, ...,<br>..., VdnDe7 | Illumina | Velvet/Oases | 1.2.10 | 23, 33, 43, 53,<br>63, 73, 83 | Default |
| PW363 | CLCdnPW1 | SOLiD | CLC *de novo* | 8.5.4 | 24 | 50 |
| PW363 | CLCdnPW2 | SOLiD | CLC *de novo* –<br>– transcript discovery<br>as reference | 9.1 | 24 | 50 |
| PW363 | VdnPW8, ...,<br>..., VdnPW10 | SOLiD | Velvet/Oases | 1.2.10 | 23, 33, 43 | Default |
| PW363 | CLCdnPW3, ...,<br>..., CLCdnPW44 | Illumina | CLC *de novo* | 8.5.4 | 21, 23, 24, 25, 30,<br>33, 35, 40, 43, 45,<br>50, 53, 55, 63 | 50, 65, 85 |
| PW363 | CLCdnPW45, ...,<br>..., CLCdnPW50 | Illumina | CLC *de novo* –<br>– transcript discovery<br>as reference | 10.0.1 | 21, 23, 33, 43,<br>53, 63 | 85 |
| PW363 | SdnPW1 | Illumina | rnaSPAdes | 3.11.1 | 43 | Default |
| PW363 | TPW | Illumina | Trinity | r2013-02-25 | 25 | NA |
| PW363 | VdnPW1, ...,<br>..., VdnPW7 | Illumina | Velvet/Oases | 1.2.10 | 23, 33, 43, 53,<br>63, 73, 83 | Default |
| Rywal | PBdnRY1 | PacBio Isoseq | Iso-Seq 3,<br>Cupcake ToFU | 2017 | NAP | NAP |
| Rywal | CLCdnRY1, ...,<br>..., CLCdnRY6 | Illumina | CLC *de novo* | 9.1 | 21, 23, 33, 43,<br>53, 63 | 85 |
| Rywal | CLCdnRY7, ...,<br>..., CLCdnRY12 | Illumina | CLC *de novo* –<br>– transcript discovery<br>as reference | 10.1.1 | 21, 23, 33, 43,<br>53, 63 | 85 |
| Rywal | SdnRY1 | Illumina | rnaSPAdes | 3.11.1 | 43 | Default |
| Rywal | VdnRY1, ...,<br>..., VdnRY7 | Illumina | Velvet/Oases | 1.2.10 | 23, 33, 43, 53,<br>63, 73, 83 | Default |

NAP – not applicable NA – not available

### Decreasing redundancy of assemblies and annotation

739 mio Désirée short reads were assembled into 3,765,661 potential transcripts, 284 mio PW363 short reads were assembled into 6,022,291 potential transcripts, and 710 mio Rywal short reads and 1.4 mio Rywal PacBio sequences were assembled into 1,912,821 potential transcripts. While generation of several transcriptomes from diverse input data and various parameter combinations increases the likelihood of capturing and accurately assembling transcripts [39], redundancy reduction without loss of information and error removal from the over-assemblies is required. All cultivar-specific transcriptome assemblies, compiled into cultivar-specific transcriptome over-assembly, were initially filtered with the tr2aacds pipeline (part of EvidentialGene v2016.07.11, [12]) which consists of four steps. First, all perfect redundant nucleotide sequences are removed using fastnrdb, part of the exonerate package [40], leaving only the transcript with the longest coding region. Next, all perfect fragments of the remaining transcripts are removed using cd-hit [41]. These first two steps are important in reducing transcriptome redundancy, as true transcripts are expected to be assembled independently by multiple of the assembly methods. Keeping the transcripts with the longest and most complete coding region helps eliminate chimeric and misassembled transcripts, as these errors tend to occur more often in UTR regions or in a manner that causes frameshifts and long, incomplete coding regions [12]).

The third and the fourth step of the tr2aacds pipeline segregate transcripts that are likely isoforms, alleles, or other variations seen at a single locus. This is done through amino acid sequence clustering, which identifies putative transcripts that vary only in silent mutations, and through reciprocal BLAST, which detects high-identity exon-sized alignments (likely isoforms). A tag is assigned to all transcripts providing detailed information on why they were discarded (e.g. perfect fragments, perfect duplicates, very high similarity, …) or why they were marked as alternatives (and which sequence they are an alternative form of). The final output of the tr2aacds pipeline are three sets of data – a non-redundant set of representative sequences (i.e. main set), a set of putative alternatives mapped to the representative set (i.e. alt set), and a discarded set (i.e. drop set) of redundant sequences. It is important to note that not all dropped sequences are of poor quality or incorrect – many of them are dropped due to full or partial redundancy.

Representative and alternative sets (termed okay sets by EvidentialGene) were merged into initial cultivar reference transcriptomes and, as tr2aacds only uses internal evidence for data curation, used in further external evidence for assembly validation, filtering and annotation steps (Figure 2). The *de novo* cultivar-specific transcript sets were first mapped to the DM reference genome by STARlong 2.6.1d [42] using parameters optimized for *de novo* transcriptome datasets (all scripts are deposited at FAIRDOMHub (fairdomhub.org) project home page, [21]). Aligned transcripts were analysed with MatchAnnot to identify transcripts that match the PGSC or ITAG gene models. Domains were assigned to the polypeptide dataset using InterProScan software package v5.37-71.0 [43]. For all transcripts and coding sequences, annotations using DIAMOND v0.9.24.125 [44] were generated by querying UniProt (www.uniprot.org) retrieved databases (E-value cut-off 10–05 and query transcript/cds and target sequence alignment coverage higher or equal to 50%; custom databases: *Solanum tuberosum*, *Solanaceae*, plants). Initially assembled transcriptomes were also screened for nucleic acid sequences that may be of vector origin (vector segment contamination) using VecScreen plus taxonomy program v.0.16 [45] against NCBI UniVec Database (ftp.ncbi.nlm.nih.gov/pub/UniVec). Potential biological and artificial contamination was identified as up to 3.3% of sequences per cultivar, if taking into account cases when potential contaminants covered less than 1% of the sequence (number of sequences with strong, moderate and weak proof of contamination as follows: 182, 547 and 10,509 for Désirée; 48, 228 and 7,877 for PW363; 169, 179 and 4,103 for Rywal). The results from MatchAnnot, InterProScan and DIAMOND were used as biological evidence in further filtering by in-house R scripts (Supplementary scripts 1). Transcripts that did not map to the genome nor had any significant hits in either InterPro (www.ebi.ac.uk/interpro) or UniProt (www.uniprot.org) were eliminated from further analysis to obtain higher reliability of constructed transcriptomes (Supplementary Table 3, Supplementary Table 4 and Supplementary Table 5). Pajek v5.08 [46], in-house scripts, and cdhit-2d from the CD-HIT package v4.6 [41] were used to re-assign post-filtering representative and alternative classes and to obtain finalised cultivar-specific transcriptomes (Supplementary scripts 2).

The whole redundancy removal procedure reduced the initial transcriptome assemblies by 18-fold for Désirée, 38-fold for Rywal, and 24-fold for PW363. Completeness of each initial *de novo* assembly to cultivar-specific transcriptome was estimated with BUSCO (Supplementary Figure 1, Supplementary Figure 2 and Supplementary Figure 3).

Individual contributions by various assembly methods were investigated in light of what contributed to the final, cleaned cultivar transcriptomes. SOLiD assemblies (Supplementary Figure 1: CLCdnDe1, CLCdnDe8, VdnDe8-10), produced by either CLC or Velvet/Oases pipelines, contributed least to transcriptomes, which can mostly be attributed to short length of the input sequences. Interestingly, increasing k-mer size in the CLC pipeline for Illumina assemblies produced more complete assemblies according to BUSCO scores and more transcripts were selected for the initial transcriptome (Supplementary Figure 1: CLCdnDe1-7, CLCdnDe9-14). On the contrary, increasing k-mer length in Velvet/Oases pipeline lead to transcripts that were less favoured by the redundancy removal procedure (Supplementary Figure 1: VdnDe1-7). The Trinity assembly was comparable to the high k-mer CLC assemblies in transcriptome contribution and BUSCO score (Supplementary Figure 1). It might seem that the PacBio Iso-Seq transcripts contributed less than expected to the cv. Rywal transcriptome (Supplementary Figure 3), however it should be noted that a considerable number of PacBio transcripts was assigned to the EvidentialGene drop set because they had perfect or near-perfect CDS identity of transcripts assembled by CLC. The EvidentialGene pipeline also prioritised CLC-assembled transcripts over PacBio transcripts because the redundancy removal algorithm reorders the near-perfect duplicates by transcript name and only retains the first transcript listed (Supplementary Table 6).

### Potato pan-transcriptome construction

While the PGSC gene model defined transcripts as well as coding sequences, the ITAG gene model defined only coding sequences. Therefore, the potato pan-transcriptome construction was conducted at the level of CDS.

Cultivar-specific representative coding sequences (57,943 of Désirée, 43,883 of PW363 and 36,336 of Rywal) were combined with coding sequences from the merged Phureja DM gene models (17,880 and 31,442 non-redundant PGSC and ITAG genes, respectively) and subjected to the cdhit-est algorithm (global sequence identity threshold 90%, alignment coverage for the shorter sequence 75%, bandwidth of alignment 51 nt and word length of 9; Supplementary scripts 3) [41] to create potato pan-trancriptome. Sequences that did not cluster using cdhit-est were separated into tetraploid and DM datasets and subjected to the cdhit-2d algorithm (local sequence identity threshold 90%, alignment coverage for the shorter sequence 45%, bandwidth of alignment 45 nt and word length of 5; Supplementary scripts 3) [41].

Sequences that are shared by the DM merged gene model and *de novo* assembled cultivar-specific transcriptomes were designated as “core” transcripts, and sequences that were assembled in only one transcriptome were designated “genotype-specific”. The total pan-transcriptome includes 96,886 representative, non-redundant transcripts and 90,618 alternative sequences (covering alternative splice forms, allelic isoforms and partial transcripts) for those loci (Figure 3, Supplementary Figure 4, Supplementary Table 7). The core subset of the pan-transcriptome contains 68,708 sequences, among which 12% are partial sequences.

**Figure 3.**
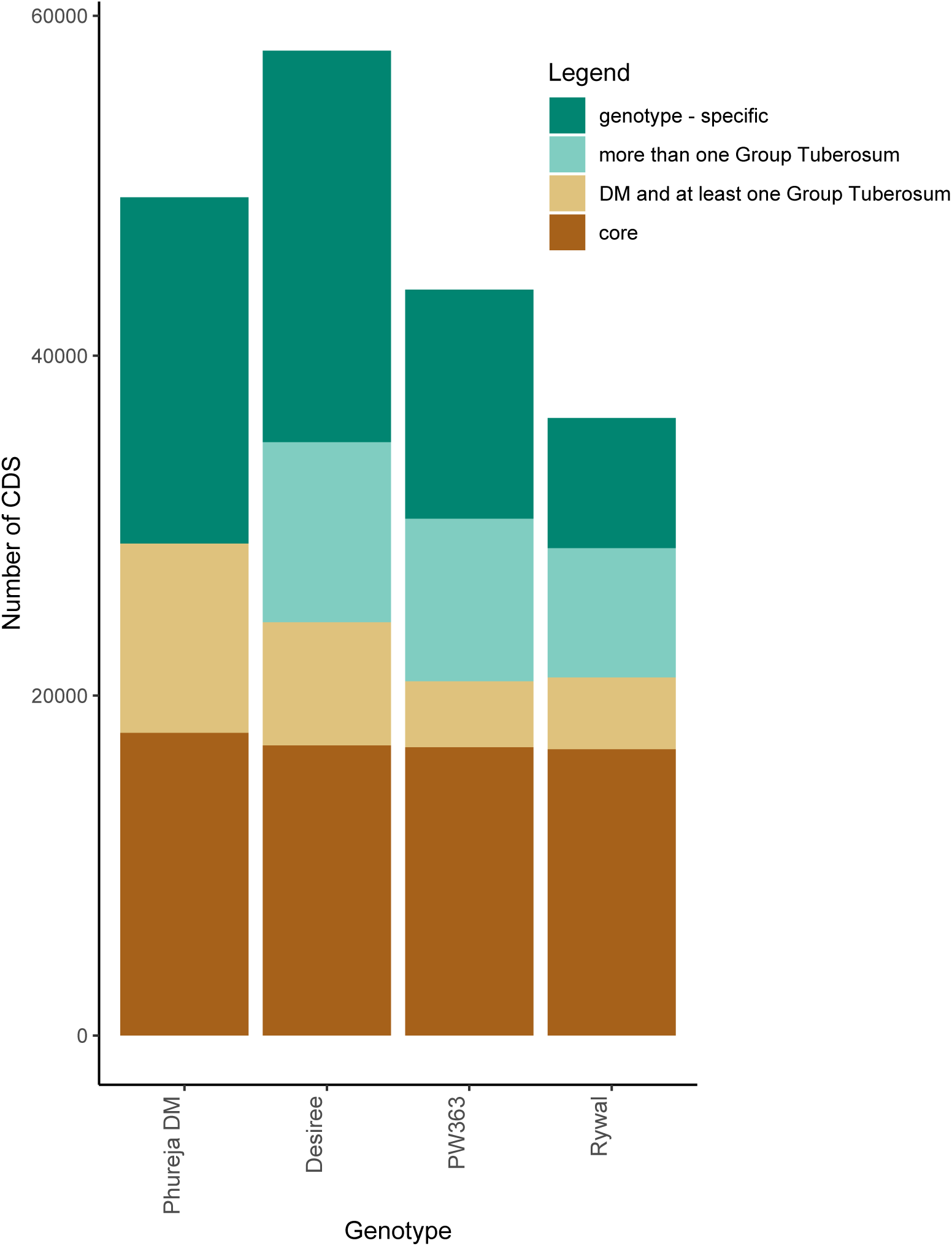
Structure of the potato pan-transcriptome. Stacked bar plot showing the overlap of paralogue groups in cultivar-specific transcriptomes and merged Phureja DM gene model. Only representative and alternative transcripts of the pan-transcriptome are counted (i.e. cultivar representative sequences) while disregarding additional cultivar alternative transcripts. For Phureja DM, the merged ITAG and PGSC DM gene models were counted. DM and at least one Group Tuberosum: sequences shared by Phureja DM and at least one tetraploid genotype, core: sequences shared among all genotypes in the pan-transcriptome.

Polyploid crop pan-genomes generally consist of many cultivar-specific genes [47]. Therefore we included all genotype-specific sequences in our potato pan-transcriptome (Figure 3, Supplementary Figure 4). This subset contains 64,529 sequences, among which 13% sequences are partial. (Supplementary Table 7). Genotype-specific transcripts are generally shorter in length than the core transcripts, however they do not differ much in the percentage of complete transcripts.

### Technical Validation

As a measure of assembly accuracy, the percentage of correctly assembled bases was obtained by mapping Illumina reads back to cultivar-specific initial transcripts using STAR v.2.6.1d RNA-seq aligner with default parameters (Table 3). To assess the quality of transcriptomes via size-based and reference-based metrics, we ran TransRate v 1.0.1 [48] on cultivar-specific transcriptomes (prior to and post filtering, Table 4), cultivar-specific representative transcript sequences and PGSC defined representative transcripts (Table 5). Comparative metrics for cultivar-specific coding sequences (CDS) were obtained using Conditional Reciprocal Best BLAST (CRBB) [49] against merged Phureja DM gene model coding sequences.

**Table 3.** Transcriptome quality control by RNA-seq reads remapping. Illumina paired-end reads used for generating assemblies were mapped back to the corresponding cultivar specific transcriptomes using STAR.

| Mapping statistics/genotype | Désirée' | PW363' | Rywal' |
| --- | --- | --- | --- |
| Number of input reads | 177,149,132 | 52,171,015 | 342,767,035 |
| Average input read length | 178 | 179 | 199 |
| **UNIQUE READS:** |  |  |  |
| Uniquely mapped reads number | 64,507,790 | 18,416,487 | 206,003,021 |
| **Uniquely mapped reads# %** | **36%** | **35%** | **60%** |
| Average mapped length | 175 | 176 | 196 |
| Number of splices*: Total | 496,170 | 267,268 | 1,700,235 |
| Number of splices*: Annotated (sjdb) | 0 | 0 | 0 |
| Number of splices*: GT/AG | 258,208 | 105,551 | 1,162,885 |
| Number of splices*: GC/AG | 10,749 | 5,693 | 79,495 |
| Number of splices*: AT/AC | 1,486 | 2,192 | 1,840 |
| Number of splices*: Non-canonical | 225,727 | 153,832 | 456,015 |
| Mismatch rate per base % | 0.50% | 0.53% | 0.59% |
| Deletion rate per base | 0.03% | 0.03% | 0.03% |
| Deletion average length | 2.72 | 2.53 | 3.02 |
| Insertion rate per base | 0.02% | 0.02% | 0.03% |
| Insertion average length | 1.93 | 1.86 | 1.91 |
| **MULTI-MAPPING READS:** |  |  |  |
| Number of reads mapped to multiple loci | 98,694,222 | 29,366,122 | 108,669,657 |
| **% of reads mapped to multiple loci#** | **56%** | **56%** | **32%** |
| Number of reads mapped to too many loci | 4,652,918 | 1,555,704 | 1,541,238 |
| **% of reads mapped to too many loci** | **2.63%** | **2.98%** | **0.45%** |
| **UNMAPPED READS:** |  |  |  |
| % of reads unmapped: too many mismatches | 0% | 0% | 0% |
| % of reads unmapped: too short | 5.25% | 5.43% | 7.75% |
| % of reads unmapped: other | 0% | 0% | 0% |
| **CHIMERIC READS:** |  |  |  |
| Number of chimeric reads | 0 | 0 | 0 |
| % of chimeric reads | 0% | 0% | 0% |

\*Number of reads crossing supposed splice sites
#Initially constructed transcriptomes (prior to filtering steps)
#relevant % of mapped reads: % of uniquely mapped reads + % of reads mapped to multiple loci

**Table 4.** Prior and post-filtering transcriptome summary statistics for potato cultivar-specific coding sequences generated by TransRate.

| TransRate metrics | Désirée |  | PW363 |  | Rywal |  |
| --- | --- | --- | --- | --- | --- | --- |
|  | Pre-filter (initial) | Post-filter | Pre-filter (initial) | Post-filter | Pre-filter (initial) | Post-filter |
| **CONTIG METRICS** |  |  |  |  |  |  |
| No. sequences | 350,271 | 197,839 | 273,216 | 159,278 | 134,755 | 79,095 |
| Sequence mean length | 504 | 792 | 516 | 775 | 459 | 707 |
| No. sequences under 200 nt | 125,465 | 25,330 | 88,230 | 17,370 | 52,653 | 13,198 |
| No. sequences over 1000 nt | 57,679 | 55,837 | 44,508 | 42,571 | 19,175 | 18,748 |
| No. sequences over 10000 nt | 23 | 23 | 3 | 3 | 1 | 1 |
| No. sequences with ORF | 132,959 | 123,546 | 109,649 | 100,222 | 43,870 | 41,523 |
| 'n90 | 369 | 444 | 366 | 429 | 351 | 390 |
| 'n50 | 1,194 | 1,209 | 1,110 | 1,131 | 1,227 | 1,218 |
| GC % | 41% | 42% | 42% | 42% | 42% | 42% |
| Ambiguous nucleotide (N) % | 0% | 0% | 0% | 0% | 0% | 0% |
| **COMPARATIVE METRICS** |  |  |  |  |  |  |
| No. seq. with CRBB hits* | 160,295 | 138,131 | 138,443 | 116,834 | 66,258 | 55,239 |
| No. reference seq. with CRBB hits* | 29,858 | 27,642 | 25,739 | 23,839 | 23,549 | 22,163 |
| coverage50#* | 25,991 | 24,586 | 21,875 | 20,620 | 20,258 | 19,538 |
| coverage95#* | 19,329 | 18,246 | 15,664 | 14,727 | 14,967 | 14,470 |
| Reference coverage* | 65% | 63% | 56% | 54% | 53% | 52% |

' the largest contig size at which at least 90% or 50% of bases are contained in contigs at least this length
\* reference-based summary statistics (merged Phureja DM coding sequences were used as reference)

**Table 5.**
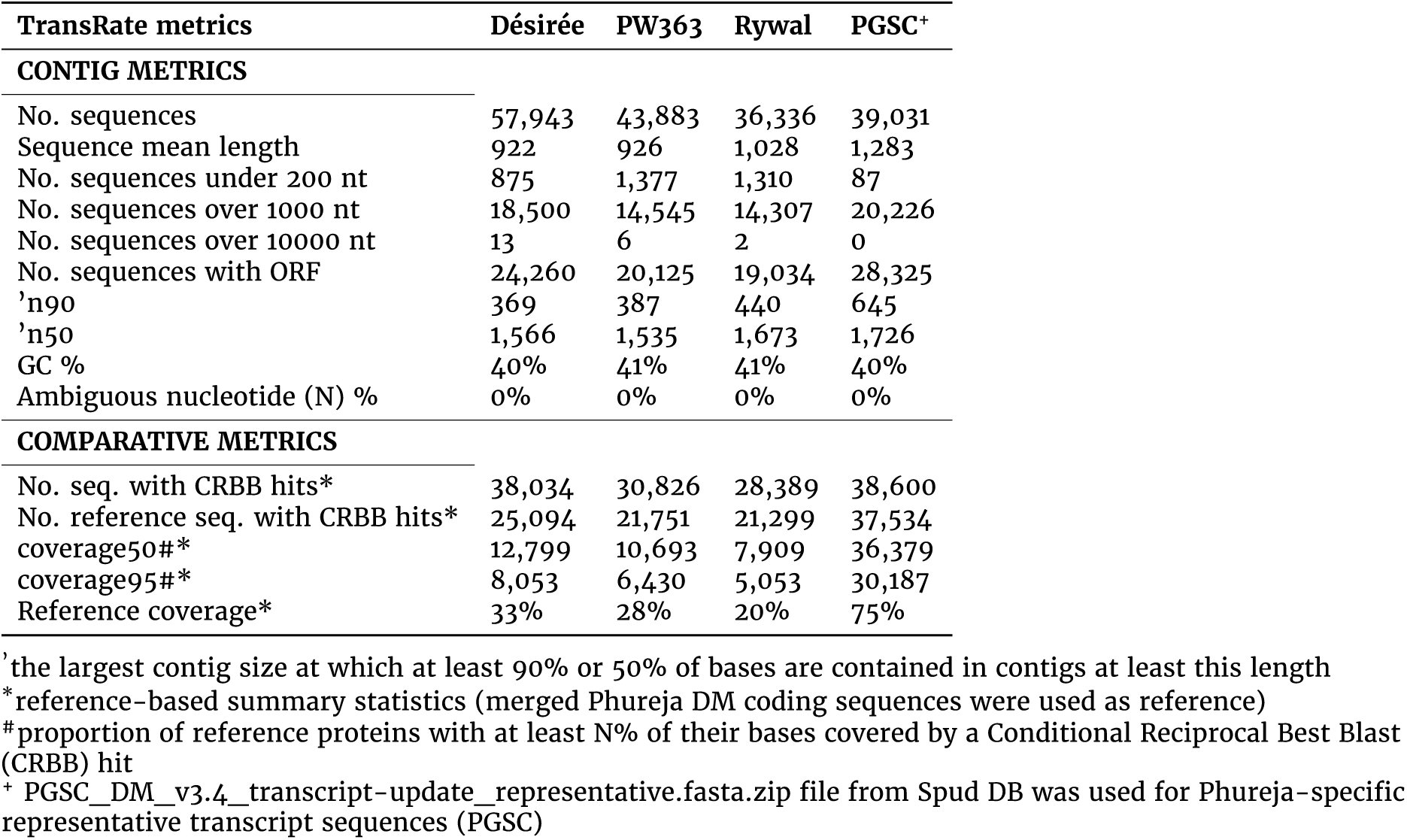
Summary statistics for potato cultivar-specific representative transcript sequences generated by TransRate.

| TransRate metrics | Désirée | PW363 | Rywal | PGSC<sup>+</sup> |
| --- | --- | --- | --- | --- |
| **CONTIG METRICS** |  |  |  |  |
| No. sequences | 57,943 | 43,883 | 36,336 | 39,031 |
| Sequence mean length | 922 | 926 | 1,028 | 1,283 |
| No. sequences under 200 nt | 875 | 1,377 | 1,310 | 87 |
| No. sequences over 1000 nt | 18,500 | 14,545 | 14,307 | 20,226 |
| No. sequences over 10000 nt | 13 | 6 | 2 | 0 |
| No. sequences with ORF | 24,260 | 20,125 | 19,034 | 28,325 |
| 'n90 | 369 | 387 | 440 | 645 |
| 'n50 | 1,566 | 1,535 | 1,673 | 1,726 |
| GC % | 40% | 41% | 41% | 40% |
| Ambiguous nucleotide (N) % | 0% | 0% | 0% | 0% |
| **COMPARATIVE METRICS** |  |  |  |  |
| No. seq. with CRBB hits* | 38,034 | 30,826 | 28,389 | 38,600 |
| No. reference seq. with CRBB hits* | 25,094 | 21,751 | 21,299 | 37,534 |
| coverage50#* | 12,799 | 10,693 | 7,909 | 36,379 |
| coverage95#* | 8,053 | 6,430 | 5,053 | 30,187 |
| Reference coverage* | 33% | 28% | 20% | 75% |

' the largest contig size at which at least 90% or 50% of bases are contained in contigs at least this length
\*reference-based summary statistics (merged Phureja DM coding sequences were used as reference)

To estimate the measure of completeness and define the duplicated fraction of assembled transcriptomes (prior and post filtering cultivar-specific transcriptomes and pan-transcriptome), BUSCO v3 [50] scores were calculated using embryophyta_odb9 [51] lineage data (Table 6). At the cultivar-specific transcriptome level, the most diverse dataset in terms of tissues and experimental conditions, the cv. Désirée dataset, resulted in the highest BUSCO score as expected. The success in classification of representative and alternative transcripts is evident from the pan-transcriptome BUSCO scores (i.e. differences in single-copy and duplicated BUSCOs for representative and alternative dataset). The highest number of fragmented BUSCOs is observed for the breeding clone PW363, which we can probably attribute to the highest number of short-contig assemblies. Furthermore, the long-read assembly presumably contributed to the shift in favour of single-copy BUSCOs (Table 6) and uniquely mapped reads (Table 3) for cv. Rywal.

**Table 6.** Assessment of completeness of constructed transcriptomes. Percentage of BUSCOs identified in each transcriptome assembly step.

| cv. Désirée | initial<br>rep+alt | post 1st filtering<br>rep+alt | final<br>rep+alt |
| --- | --- | --- | --- |
| (S) | 37.8 | 37.8 | 37.4 |
| (D) | 59.4 | 59.2 | 58.4 |
| (C) | **97.2** | **97.0** | **95.8** |
| (F) | 1.1 | 1.2 | 1.4 |
| (M) | 1.7 | 1.8 | 2.8 |
| breeding clone PW363 | initial<br>rep+alt | post 1st filtering<br>rep+alt | final<br>rep+alt |
| (S) | 39.9 | 39.2 | 38.4 |
| (D) | 51.7 | 51.2 | 50.9 |
| (C) | **91.6** | **90.4** | **89.3** |
| (F) | 2.9 | 3.4 | 3.5 |
| (M) | 5.6 | 6.2 | 7.2 |
| cv. Rywal | initial<br>rep+alt | post 1st filtering<br>rep+alt | final<br>rep+alt |
| (S) | 55.8 | 55.8 | 55.1 |
| (D) | 35.2 | 34.8 | 34.7 |
| (C) | **91.0** | **90.6** | **89.8** |
| (F) | 2.4 | 2.6 | 2.7 |
| (M) | 6.5 | 6.9 | 7.5 |
| pan-transcriptome | rep | alt | rep+alt |
| (S) | 92.2 | 11.0 | 3.9 |
| (D) | 6.1 | 85.9 | 95.6 |
| (C) | **98.3** | **96.9** | **99.4** |
| (F) | 1.4 | 1.3 | 0.3 |
| (M) | 0.3 | 1.7 | 0.3 |

(S): Complete and single-copy BUSCOs %;
(D): Complete and duplicated BUSCOs %
(C): Complete BUSCOs (S + D) %
(F): Fragmented BUSCOs %
(M): Missing BUSCOs %
rep: representative
alt: alternative
\*Database size: 1440

To inspect the quality of paralogue cluster assignments, multiple sequence alignments (MSA) using MAFFT v7.271 [52] were conducted on representative and alternative sequences from paralogue clusters (Supplementary Table 7) containing sequences from each of the four genotypes (Désirée, PW363, Rywal and DM). Alignments were visualized using MView v1.66 [53] (Supplementary File 2). These alignments were used to quality check *de novo* transcriptome assemblies and helped us optimise the pipeline. The alignments within groups showed differences that can be attributed to biological diversity, e.g. SNPs and INDELS as well as alternative splicing. However, we advise the users to check the available MSA for their gene of interest as some miss-assemblies might still occur (Supplementary File 2, Supplementary scripts 4).

To estimate the proportion of transcripts originating from organisms other than potato, we performed a taxonomic classification of all cultivar-specific transcriptome sequences using Centrifuge v1.0.4 [54] and the NCBI nt database. We used pavian [55] to generate classification summary reports (Supplementary File 3). The transcriptomes include altogether less than 1% bacterial, viral, fungal and protozoan transcripts. Other non-plant sequences are from common plant pests such the arachnid *Tetranychus urticae* in PW363 transcriptome (cca. 4%) and insects *Leptinotarsa decemlineata* (cca. 0.1%) and *Myzus persicae* (cca. 0.1%) in the Désirée transcriptome.

### Sanger sequencing confirmation of assembled transcripts

To further validate the quality of the constructed Rywal reference transcriptome, the cultivar-specific assembled transcript coding for NPR1-1 protein in cv. Rywal was compared to sequences amplified from isolated cDNA and sequenced by Sanger method. Total RNA was isolated from 4-week-old cv. Rywal plants using RNeasy Plant Mini kit (Qiagen). Residual genomic DNA was digested with DNase I (Qiagen). Treated total RNA was reversed transcribed using High Capacity cDNA Reverse Transcription Kit (Applied Biosystems). The full-length CDS of NPR1-1 (Sotub07g016890.1.1, ITAG genome annotation) was amplified from the cv. Rywal cDNA with forward (5’-ATGGAGAGCGGCCACGAGA-3’) and reverse (5’-CTACTTTTTTCTAAACTTGTGACTGACATT-3’) primers. The PCR product was inserted into pJET1.2/blunt vector with CloneJET PCR Cloning Kit (Thermo Scientific) and introduced into OneShot TOP10 Chemically Competent *Escherichia coli* cells. The plasmids were isolated from 8 transformed colonies, grown on selection media, using the GenElute Plasmid Miniprep Kit (Sigma-Aldrich). Inserts were sequenced (GATC Services, Eurofins Genomics) using pJET1.2 sequencing primers (Thermo Scientific) as well as forward (5’-CTCCAAGGTTGTGAACGAGGTACTT-3’) and reverse (5’-AAGTACCTCGTTCACAACCTTGGAG-3’) insert-specific primers, designed to ensure full sequence coverage in both directions.

A multiple sequence alignment comparing Sanger sequences with NPR1-1 coding sequence from the assembled cv. Rywal transcriptome (paralogue cluster stCuSTr-R_29366) and Phureja DM gene model (Sotub07g016890.1.1) was constructed using Geneious Prime 2020.1.1 (https://www.geneious.com) [56]. The Sanger sequencing revealed the presence of several distinct gene variants in the analysed colonies, differing in the SNP pattern. The PBdnRY1_9437 sequence is validated by complete sequence identity with the two colonies (seq7 and 8), while the CLCdnRY_10265 shares two SNPs with seq4 - 6, matching the SNP pattern partially. Although the Phureja DM gene model also shares two SNPs with some of the colonies, its overall SNP pattern differs significantly from cv. Rywal, distinguishing the cultivar-specific transcripts from that of Phureja DM (Figure 4).

**Figure 4.**
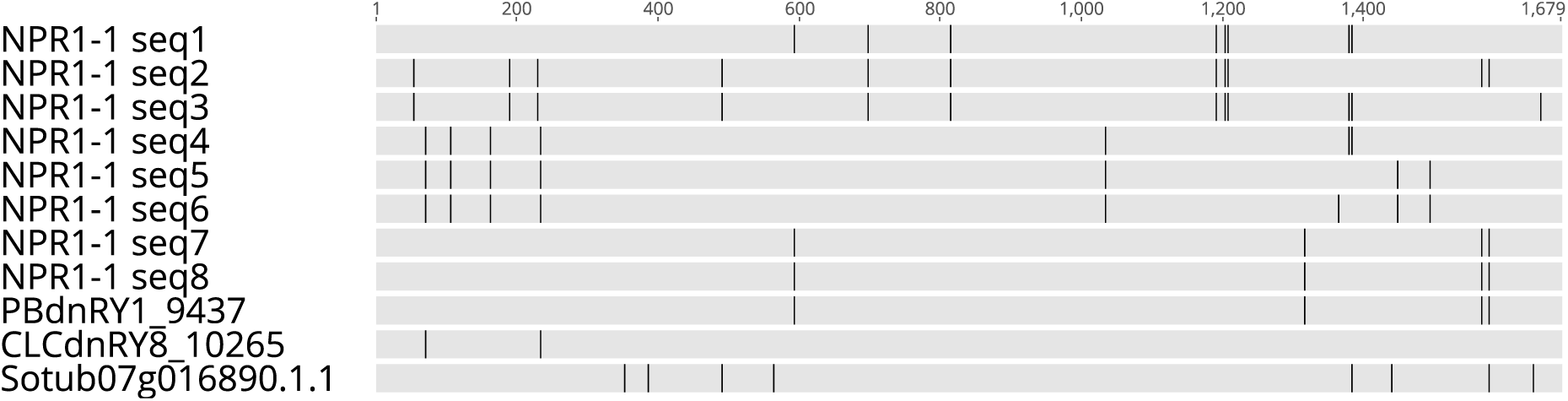
Sanger sequencing validates the constructed cultivar specific transcriptome. Multiple sequence alignment of NPR1-1 coding sequence obtained from eight *E. coli* colonies (NPR1-1 seq1 - 8) by the Sanger method, assembled short or long-read cv. Rywal transcripts and Phureja DM gene model (Sotub07g016890.1.1). Grey - sequence identity, black - SNPs. The alignment was prepared and visualised with Geneious Prime 2020.1.1. [56]

## Usage Notes

### Insights into variability of potato transcriptomes

Based on the comparison of cultivar-specific transcriptomes we identified cca. 23,000, 13,000, and 7,500 paralogue groups of transcripts in cv. Désirée, breeding clone PW363 and cv. Rywal, respectively, not present in the merged Phureja DM gene model. The addition of Iso-Seq dataset in the case of cv. Rywal confirms that long reads contribute to less fragmentation of the *de novo* transcriptome. It is therefore recommended to generate at least a subset of data with one of the long-read technologies to complement the short-read RNA-seq. As it can be seen by the reduction rate in PW363 (24-fold), producing additional short-read assemblies does not contribute to the transcriptome quality as much as having several tissues or a combination of 2*nd* and 3*rd* generation sequencing (38-fold Rywal).

From all four genotypes, cv. Désirée has the highest number of cultivar-specific representative transcripts, which can be attributed to having the most diverse input dataset used for the *de novo* assemblies in terms of tissues sequenced (stem, seedlings and roots) and experimental conditions covered. Cv. Désirée also benefited from the inclusion of a DSN Illumina library to capture low level expressed transcripts. However, even the leaf-specific reference transcriptomes of cv. Rywal and breeding clone PW363 include thousands of specific genes, indicating that cultivar specific gene content is common. Remarkably, we identified several interesting features when inspecting paralogue groups of transcripts, demonstrating the variability of sequences in potato haplotypes and the presence of the alternative splicing variants that contribute to the pan-transcriptome (Figure 5, Supplementary File 2).

**Figure 5.**
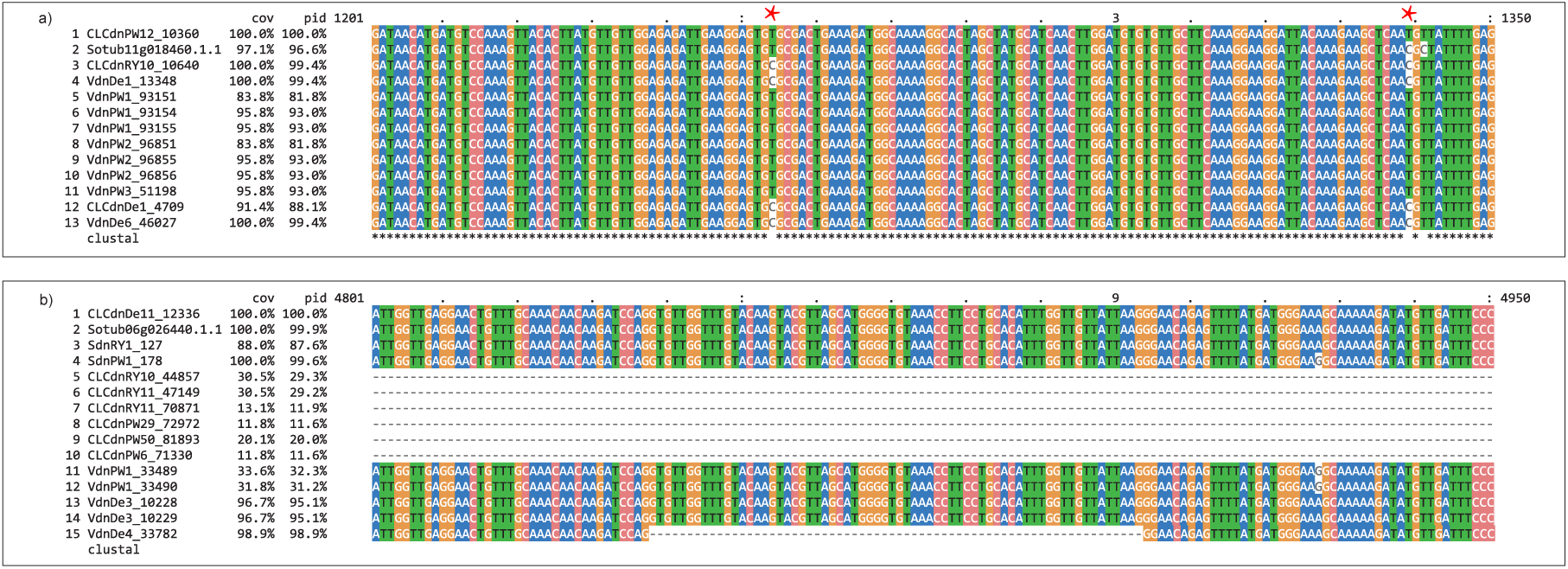
Transcript variants present in pan-transcriptome paralogue gene groups. a) Alignment part of stPanTr_010101 with two PW363-specific SNPs marked by red dots. Such SNPs can be used to design cultivar- or allele-specific qPCR assays. b) Alignment part of stPanTr_074336 showing an alternative splice variant in Désirée, (VdnDe4_33782). Both multiple sequence alignments were made using ClustalOmega v 1.2.1 [59] and visualized with MView v 1.66 [53]. The remaining alignments can be found in Supplementary File 2.

It should be noted, that the reconstructed transcriptomes include also the meta-transcriptome stemming from microbial communities present in sampled potato tissues (Supplementary File 3). We decided not to apply any filter on these transcripts. Inclusion of meta-transcripts makes it possible to also investigate the diversity of plant-associated endo- and epiphytes. The majority of these microbial transcripts will have microbial annotations, facilitating their future removal when necessary for other experiments.

### Cultivar-specific transcriptomes can improve high-throughput sequencing analyses

Most gene expression studies have been based on either potato UniGenes, assembled from a variety of potato expressed sequence tags (e.g. StGI, POCI), or the reference DM genome transcript models. Studies based on any of these resources have provided useful information on potato gene expression, but also have major drawbacks.

When using the DM genome as a reference for mapping RNA-Seq reads, the potato research community faces the existence of two overlapping, but not identical, gene model predictions. When using either of available GFFs, we are missing some of the genes known to be encoded in the assembled scaffold. The newly generated merged DM-based GTF helps to circumvent this problem, but even in the merged GTF the cultivar-specific genes and variations are not considered. Differences in expression and important marker transcripts can therefore be missed. In addition, the computational prediction of DM transcript isoforms is incomplete and, in some cases, gene models are incorrectly predicted. On the other hand, the inherent heterogeneity and redundancy of UniGenes or similarly combined transcript sets causes the mapping of short reads to multiple transcripts, thus making the result interpretation more difficult. The cultivar-specific transcriptomes presented here are an improvement, as they include several transcripts missing in the Phureja DM transcriptome. The transcriptomes are also a valuable asset for other high-throughput sequencing applications, such as sRNA-Seq, Degradome-Seq or ATAC-Seq, as we now have more detailed information also on transcript variability within one locus.

The benefit of using cultivar-specific transcriptomes was demonstrated through mapping statistics for Désirée leaf samples under drought stress [57] to Désirée, ITAG/PGSC merged and PGSC representative transcriptome sequences. Taking all three samples together, 5.5% more reads were uniquely mapped to Désirée than the ITAG/PGSC Phureja DM gene models (Table 7). From the reads mapped to Désirée transcriptome, 5.3% mapped to Désirée-specific transcripts and 8.2% to transcripts specific to Group Tuberosum genotypes (Supplementary Table 8).

**Table 7.** Mapping of independent dataset to newly assembled cultivar specific reference transcriptome. Mapping statistics for Désirée leaf samples under drought stress to Désirée, ITAG/PGSC merged and PGSC representative transcriptome sequences is shown.

| Reference | Désirée | ITAG/PGSC | PGSC | Désirée | ITAG/PGSC | PGSC |
| --- | --- | --- | --- | --- | --- | --- |
| Mapping statistics/Sample | SRR10416847 |  |  | SRR10416848 |  |  |
| Number of input reads | 14,953,659 |  |  | 14,610,172 |  |  |
| Average input read length | 252 |  |  | 252 |  |  |
| UNIQUE READS: |  |  |  |  |  |  |
| Uniquely mapped reads# % | 74% | 69% | 71% | 73% | 67% | 69% |
| Average mapped length | 246 | 244 | 246 | 246 | 244 | 246 |
| Number of splices: Total | 123,490 | 104,658 | 159,842 | 127,414 | 110,423 | 165,595 |
| Number of splices: Non-canonical | 40,960 | 43,764 | 45,750 | 40,898 | 45,459 | 42,978 |
| Mismatch rate per base % | 0.75% | 0.94% | 1.00% | 0.74% | 0.93% | 1.00% |
| Deletion rate per base | 0.05% | 0.04% | 0.06% | 0.05% | 0.04% | 0.06% |
| Deletion average length | 3.42 | 3.46 | 2.88 | 3.41 | 3.47 | 2.89 |
| Insertion rate per base | 0.03% | 0.02% | 0.04% | 0.03% | 0.02% | 0.04% |
| Insertion average length | 2.21 | 2.99 | 2.52 | 2.24 | 3.00 | 2.53 |
| MULTI-MAPPING READS: |  |  |  |  |  |  |
| % of reads mapped to multiple loci# | 4.94% | 1.95% | 2.74% | 4.86% | 1.68% | 2.52% |
| % of reads mapped to too many loci | 0% | 0% | 0% | 0% | 0% | 0% |
| UNMAPPED READS: |  |  |  |  |  |  |
| % of reads unmapped: too many mismatches | 0% | 0% | 0% | 0% | 0% | 0% |
| % of reads unmapped: too short | 21% | 29% | 27% | 22% | 31% | 29% |
| % of reads unmapped: other | 0% | 0% | 0% | 0% | 0% | 0% |
| Mapping statistics/Sample | SRR10416849 |  |  | all samples |  |  |
| Number of input reads | 14,755,430 |  |  | 44,319,261 |  |  |
| Average input read length | 252 |  |  | 252 |  |  |
| UNIQUE READS: |  |  |  |  |  |  |
| Uniquely mapped reads# % | 49% | 44% | 46% | 66% | 60% | 62% |
| Average mapped length | 245 | 243 | 246 | 246 | 244 | 246 |
| Number of splices: Total | 95,409 | 77,103 | 115,083 | 346,313 | 292,184 | 440,520 |
| Number of splices: Non-canonical | 33,269 | 31,224 | 31,065 | 115,127 | 120,447 | 119,793 |
| Mismatch rate per base % | 0.75% | 0.94% | 1.01% | 0.75% | 0.94% | 1.00% |
| Deletion rate per base | 0.06% | 0.05% | 0.07% | 0.05% | 0.05% | 0.06% |
| Deletion average length | 3.23 | 3.25 | 2.83 | 3.36 | 3.41 | 2.87 |
| Insertion rate per base | 0.03% | 0.03% | 0.04% | 0.03% | 0.02% | 0.04% |
| Insertion average length | 2.27 | 3.15 | 2.56 | 2.24 | 3.04 | 2.54 |
| MULTI-MAPPING READS: |  |  |  |  |  |  |
| % of reads mapped to multiple loci# | 3.60% | 1.19% | 1.94% | 4.47% | 1.61% | 2.40% |
| % of reads mapped to too many loci | 0% | 0% | 0% | 0% | 0% | 0% |
| UNMAPPED READS: |  |  |  |  |  |  |
| % of reads unmapped: too many mismatches | 0% | 0% | 0% | 0% | 0% | 0% |
| % of reads unmapped: too short | 47% | 55% | 52% | 30% | 38% | 36% |
| % of reads unmapped: other | 0% | 0% | 0% | 0% | 0% | 0% |

RNA-seq data from Désirée leaf samples under drought stress retrieved from the GEO Series GSE14,0083 - "Transcriptome profiles of contrasting potato (*Solanum tuberosum* L.) genotypes under water stress".
No chimeric reads detected.

Cultivar-specific transcriptomes may also help improve mass-spectroscopy based proteomics. Comprehensive protein databases, obtained from transcriptomic data, offer a higher chance of finding significant targets with peptide spectrum match algorithms, thus enhancing the detection and sensitivity of protein abundance measurements [58].

### Using transcriptomes to inform qPCR amplicon design

Aligning transcript coding sequences from a StPanTr paralogue cluster can be used to inform qPCR primer design in order to study gene expression in different cultivars or specific isoforms by selecting variable transcript regions (Figure 5). On the other hand, when qPCR assays need to cover multiple cultivars, the nucleotide alignments can be inspected for conservative regions for design.

### Code availability

All used Bash, Perl, Python, and R/Markdown custom code and scripts complemented with intermediate and processed data (input and output files), and all other supporting information that enable reproduction and re-use are available at FAIRDOMHub under project name _p_stRT (fairdomhub.org/projects/161) under CC BY 4.0 licenses. Data were locally stored in a ISA-tab compliant project directory tree generated using pISA-tree (github.com/NIB-SI/pISA) and uploaded to FAIRDOMHub repository using FAIRDOMHub API and R package pisar (github.com/NIB-SI/pisar). Scripts and files referenced in the text are listed in the subsection Supplementary Information.

## Acknowledgements

We thank Robin Buell for potato PGSC-ITAG gene model reference table, Thomas Doak and Don Gilbert for advice on using EvidentialGene scripts, Henrik Krnec for BLAST output parser, Andrej Blejec for FAIRDOMhub API usage in R, Špela Baebler for providing DSN Illumina-sequenced samples and Karmen Pogačar for providing Rywal cDNA samples for technical validation.

## Author contributions

M.P., K.G., Ž.R. and M. Zagorščak participated in study design and evaluation of transcriptomes. A.C. provided Rywal Illumina-sequenced samples. M.P. collected and pre-processed RNA-Seq datasets and produced CLC, Velvet/Oases and rnaSPAdes *de novo* assemblies. M.Zouine produced Trinity assemblies. E.T. processed Iso-Seq data. M.P. and M.Zagorščak run tr2aacds scripts and transcriptome annotation, filtering and evaluation software. Ž.R. merged PGSC and ITAG gene models of reference potato genome. M.P., Ž.R. and M.Zagorščak generated the pan-transcriptome. Š.T. conducted bacterial transformation and Sanger sequencing analysis. K.G. secured funding, and managed the project. M.P., M.Zagorščak, Ž.R., Š.T. and K.G. wrote and edited the manuscript. S.S. helped with interpretation of EvidentialGene results, provided advice on filtering of transcriptomes and language editing. All authors have read and commented the manuscript and approved the final submission.

| Contributor Role | Authors |
| --- | --- |
| Conceptualization | K.G., M.P., Ž.R., M. Zagorščak |
| Supervision | K.G. |
| Project Administration | K.G. |
| Investigation | M.P., M. Zagorščak |
| Formal Analysis | M.P., Ž.R., M. Zagorščak |
| Software | M.P., Ž.R., E.T., M. Zagorščak |
| Methodology | K.G., M.P., Ž.R., M. Zagorščak |
| Validation | M.P., Ž.R., S.S., Š.T., M. Zagorščak |
| Data Curation | M.P., M. Zagorščak |
| Resources | A.C., K.G., M.P., E.T., Š.T., M. Zouine |
| Funding Acquisition | K.G. |
| Writing – Original Draft Preparation | M.P., M. Zagorščak |
| Writing – Review & Editing | all authors |
| Visualization | M.P., M.Zagorščak |

## Supplementary Information

### Supplementary tables

1. Supplementary Table 1 - **Detailed sample information table used to generate the *de novo* transcriptome assemblies**. Raw and processed reads summary. Layer _p_stRT/_I_STRT/ _S_01_sequences/reports/, http://doi.org/10.15490/FAIRDOMHUB.1.DATAFILE.3090.1
2. Supplementary Table 2 - **Detailed *de novo* assemblies information table**. Primary potato transcriptome assemblies summary listing parameters used for short-read *de novo* assembly generation. Layer _p_stRT/_I_STRT/_S_02_denovo/reports/, http://doi.org/10.15490/FAIRDOMHUB.1.DATAFILE.3091.1
3. Supplementary Table 3 - **Désirée biological evidence filtering results**. Output of 1*st* filtering step by biological evidence for cv. Désirée. Layer _p_stRT/_I_STRT/_S_03_stCuSTr/_A_03.1_filtering/reports/, http://doi.org/10.15490/FAIRDOMHUB.1.DATAFILE.3407.1
4. Supplementary Table 4 - **PW363 biological evidence filtering results**. Output of 1*st* filtering step by biological evidence for breeding clone PW363. Layer _p_stRT/ _I_STRT/_S_03_stCuSTr/_A_03.1_filtering/reports/, http://doi.org/10.15490/FAIRDOMHUB.1.DATAFILE.3406.1
5. Supplementary Table 5 - **Rywal biological evidence filtering results**. Output of 1*st* filtering step by biological evidence for cv. Rywal. Layer _p_stRT/_I_STRT/_S_03_stCuSTr/_A_03.1_filtering/reports/, http://doi.org/10.15490/FAIRDOMHUB.1.DATAFILE.3405.1
6. Supplementary Table 6 - **EvidentialGene Summary Statistics for PacBio sequences.** Layer _p_stRT/_I_STRT/_S_03_stCuSTr/_A_02.2_assembly-contribution-count/reports/, http://doi.org/10.15490/FAIRDOMHUB.1.DATAFILE.3363.1
7. Supplementary Table 7 - Paralogue cluster information for cultivar-specific and pan-transcriptome sequences extended with annotations and quality classification. Layer _p_stRT/_I_STRT/_S_04_stPanTr/_A_02_cdhit_3cvs-GFFmerged/reports/, http://doi.org/10.15490/FAIRDOMHUB.1.DATAFILE.3721.2
8. Supplementary Table 8 - Read count summary for Désirée drought samples mapped to the representative Phureja DM and Désirée reference transcriptomes. Layer _p_stRT/_I_STRT/_S_04_stPanTr/_A_07_Desiree-mapping/reports/, http://doi.org/10.15490/fairdomhub.1.datafile.3722.2

### Supplementary figures

1. Supplementary Figure 1 - **Contribution of individual *de novo* assemblies to cultivar Désirée transcriptome.** Proportion of all contigs in *de novo* assembly (blue bars) and proportion of EvidentialGene okay set (green bars), and the number of complete BUSCOs (dots) using embryophyta_odb9 set are shown. Assembly software abbreviations: CLCdn - CLC Genomics Workbench, Vdn - Velvet. Layer _p_stRT/_I_STRT/_S_03_stCuSTr/_A_02.2_assembly-contribution-count/reports/, http://doi.org/10.15490/FAIRDOMHUB.1.DATAFILE.3107.1
2. Supplementary Figure 2 - **Contribution of individual *de novo* assemblies to cultivar PW363 transcriptome.** Proportion of all contigs in *de novo* assembly (blue bars) and proportion of EvidentialGene okay set (green bars), and the number of complete BUSCOs (dots) using embryophyta_odb9 set are shown. Assembly software abbreviations: CLCdn - CLC Genomics Workbench, Vdn - Velvet, Sdn - SPAdes. Layer _p_stRT/_I_STRT/_S_03_stCuSTr/_A_02.2_assembly-contribution-count/reports/, http://doi.org/10.15490/FAIRDOMHUB.1.DATAFILE.3108.1
3. Supplementary Figure 3 - **Contribution of individual *de novo* assemblies to cultivar Rywal transcriptome.** Proportion of all contigs in *de novo* assembly (blue bars) and proportion of EvidentialGene okay set (green bars), and the number of complete BUSCOs (dots) using embryophyta_odb9 set are shown. Assembly software abbreviations: CLCdn - CLC Genomics Workbench, Vdn - Velvet, Sdn - SPAdes, PBdn - PacBio. Layer _p_stRT/_I_STRT/_S_03_stCuSTr/_A_02.2_assembly-contribution-count/reports/, http://doi.org/10.15490/FAIRDOMHUB.1.DATAFILE.3109.1
4. Supplementary Figure 4 - **Venn diagrams showing the overlap of paralogue clusters in cultivar-specific transcriptomes and merged Phureja DM gene models**. Representatives and alternatives of the stPanTr (pan-transcriptome) paralogue cluster are counted as well as alternatives defined at stCuSTr (cultivar-specific transcriptome) step. For Phureja, the merged ITAG and PGSC DM gene models were counted. Layer _p_stRT/_I_STRT/_S_04_stPanTr/_A_02_cdhit_3cvs-GFFmerged/reports/, http://doi.org/10.15490/FAIRDOMHUB.1.DATAFILE.3409.2

### Supplementary files

1. Supplementary File 1 - **Merged ITAG/PGSC Phureja DM gene models**. Archive containing GTF, CDS and peptide fasta files for merged ITAG and PGSC gene models for *S. tuberosum* Group Phureja DM genome v4.04 http://doi.org/10.15490/FAIRDOMHUB.1.DATAFILE.3408.1
2. Supplementary File 2 - **Multiple sequence alignments using ClustalOmega v1.2.1 or MAFFT v7.271 and MView v1.66**. Paralogue cluster on representative and alternative sequences, at least one from each of the four genotypes. Layer _p_stRT/_I_STRT/_S_04_stPanTr/_A_05_MSA_3cvs-GFFmerged/reports/, http://doi.org/10.15490/FAIRDOMHUB.1.DATAFILE.3691.1
3. Supplementary File 3 - **Evaluation of constructed reference transcriptomes for presence of the metagenome.** Pavian visualisation of the Centrifuge taxonomic classification program output forDésirée, PW363 and Rywal representative transcripts is shown. Layer _p_stRT/_I_STRT/_S_04_stPanTr/_A_08_cdhit_centrifuge_3cvs-GFFmerged/reports/, http://doi.org/10.15490/FAIRDOMHUB.1.DATAFILE.3509.1

## Supplementary scripts

1. Supplementary scripts 1 - **Evidence filtering**. Corresponding scripts for biological evidence filtering step. Layer _p_stRT/_I_STRT/_S_03_stCuSTr/_A_03.1_filtering/scripts/, http://doi.org/10.15490/FAIRDOMHUB.1.ASSAY.1170.1
2. Supplementary scripts 2 - **stCuSTr paralogue clustering**. Corresponding scripts for StCuSTr post-filtering representative (non-redundant) and alternative classes reassignment step. Layer _p_stRT/ _I_STRT/_S_03_stCuSTr/_A_03.2_components/scripts/, http://doi.org/10.15490/FAIRDOMHUB.1.ASSAY.1171.1
3. Supplementary scripts 3 - **stPanTr paralogue clustering**. Corresponding scripts for StPanTr representative and alternative transcripts classification step. Layer _p_stRT/_I_STRT/_S_04_stPanTr/_A_02_cdhit_3cvs-GFFmerged/scripts/, http://doi.org/10.15490/FAIRDOMHUB.1.ASSAY.1263.2
4. Supplementary scripts 4 - **MSA**. Corresponding scripts for MSA step. Layer _p_stRT/ _I_STRT/_S_04_stPanTr/_A_05_MSA_3cvs-GFFmerged/scripts/, http://doi.org/10.15490/FAIRDOMHUB.1.ASSAY.1270.2

## References

[1] Hardigan, M. A. et al. Genome diversity of tuber-bearing Solanum uncovers complex evolutionary history and targets of domestication in the cultivated potato. Proc. Natl. Acad. Sci. USA 114, E9999–E10008 (2017).

[2] Gan, X. et al. Multiple reference genomes and transcriptomes for Arabidopsis thaliana. Nature 477, 419–423 (2011).

[3] Hirsch, C. N. et al. Insights into the maize pan-genome and pan-transcriptome. Plant Cell 26, 121–135 (2014).

[4] Jin, M. et al. Maize pan-transcriptome provides novel insights into genome complexity and quantitative trait variation. Sci. Rep. 6, 18936 (2016).

[5] Zhao, Q. et al. Pan-genome analysis highlights the extent of genomic variation in cultivated and wild rice. Nat. Genet. 50, 278–284 (2018).

[6] Montenegro, J. D. et al. The pangenome of hexaploid bread wheat. Plant J. 90, 1007–1013 (2017).

[7] Li, Y. H. et al. De novo assembly of soybean wild relatives for pan-genome analysis of diversity and agronomic traits. Nat. Biotechnol. 32, 1045–1052 (2014).

[8] Xu, X. et al. Genome sequence and analysis of the tuber crop potato. Nature 475, 189–195 (2011).

[9] Liu, Y. et al. Comparative transcriptome analysis of white and purple potato to identify genes involved in anthocyanin biosynthesis. PLoS One 10, e0129148 (2015).

[10] Sato, S. et al. The tomato genome sequence provides insights into fleshy fruit evolution. Nature 485, 635–641 (2012).

[11] Hölzer, M. & Marz, M. De novo transcriptome assembly: A comprehensive cross-species comparison of short-read RNA-Seq assemblers. GigaScience 8, giz039 (2019).

[12] Gilbert, D. G. Genes of the pig, Sus scrofa, reconstructed with EvidentialGene. PeerJ 7, e6374 (2019).

[13] NCBI Sequence Read Archive (SRA) https://identifiers.org/ncbi/insdc.sra:SRP220411 (2019).

[14] NCBI Sequence Read Archive (SRA) https://identifiers.org/ncbi/insdc.sra:SRP040682 (2015).

[15] NCBI Sequence Read Archive (SRA) https://identifiers.org/ncbi/insdc.sra:SRP220250 (2019).

[16] NCBI Sequence Read Archive (SRA) https://identifiers.org/ncbi/insdc.sra:SRP220356 (2019).

[17] NCBI Sequence Read Archive (SRA) https://identifiers.org/ncbi/insdc.sra:SRP172523 (2019).

[18] NCBI Sequence Read Archive (SRA) https://identifiers.org/ncbi/insdc.sra:SRP069961 (2016).

[19] NCBI Sequence Read Archive (SRA) https://identifiers.org/ncbi/insdc.sra:SRP083083 (2017).

[20] NCBI Sequence Read Archive (SRA) https://identifiers.org/ncbi/insdc.sra:ERP003480 (2014).

[21] Wolstencroft, K. et al. FAIRDOMHub: a repository and collaboration environment for sharing systems biology research. Nucleic Acids Res. 45, D404–D407 (2017).

[22] GenBank https://identifiers.org/ncbi/insdc:MT210578 (2020).

[23] GenBank https://identifiers.org/ncbi/insdc:MT210579 (2020).

[24] GenBank https://identifiers.org/ncbi/insdc:MT210580 (2020).

[25] GenBank https://identifiers.org/ncbi/insdc:MT210581 (2020).

[26] GenBank https://identifiers.org/ncbi/insdc:MT210582 (2020).

[27] GenBank https://identifiers.org/ncbi/insdc:MT210583 (2020).

[28] GenBank https://identifiers.org/ncbi/insdc:MT210584 (2020).

[29] GenBank https://identifiers.org/ncbi/insdc:MT210585 (2020).

[30] FAIRDOMHub http://doi.org/10.15490/FAIRD0MHUB.1.DATAFILE.3408.1 (2020).

[31] FAIRDOMHub http://doi.org/10.15490/FAIRD0MHUB.1.ASSAY.1268.2 (2020).

[32] Hirsch, C. D. et al. Spud DB: A resource for mining sequences, genotypes, and phenotypes to accelerate potato breeding. Plant Genome 7, 1–12 (2014).

[33] Zerbino, D. R. Using the Velvet de novo assembler for short-read sequencing technologies. Curr. Protoc. in Bioinformatics 31, 11.5.1–11.5.12 (2010).

[34] Crusoe, M. R. et al. The khmer software package: enabling efficient nucleotide sequence analysis. F1000Res. 4, 900 (2015).

[35] Tseng, E. cdna_cupcake v9.0.1 (2019).

[36] Grabherr, M. G. et al. Full-length transcriptome assembly from RNA-Seq data without a reference genome. Nat. Biotechnol. 29, 644–652 (2011).

[37] Schulz, M. H., Zerbino, D. R., Vingron, M. & Birney, E. Oases: Robust de novo RNA-seq assembly across the dynamic range of expression levels. Bioinformatics 28, 1086–1092 (2012).

[38] Bushmanova, E., Antipov, D., Lapidus, A. & Prjibelski, A. D. rnaSPAdes: a de novo transcriptome assembler and its application to RNA-Seq data. GigaScience 8, giz100 (2019).

[39] He, B. et al. Optimal assembly strategies of transcriptome related to ploidies of eukaryotic organisms. BMC Genomics 16, 65 (2015).

[40] Slater, G. S. C. & Birney, E. Automated generation of heuristics for biological sequence comparison. BMC Bioinformatics 6, 31 (2005).

[41] Fu, L., Niu, B., Zhu, Z., Wu, S. & Li, W. CD-HIT: accelerated for clustering the next-generation sequencing data. Bioinformatics 28, 3150–3152 (2012).

[42] Dobin, A. et al. STAR: Ultrafast universal RNA-seq aligner. Bioinformatics 29, 15–21 (2013).

[43] Jones, P. et al. InterProScan 5: Genome-scale protein function classification. Bioinformatics 30, 1236–1240 (2014).

[44] Buchfink, B., Xie, C. & Huson, D. H. Fast and sensitive protein alignment using DIAMOND. Nat. Methods 12, 59–60 (2014).

[45] Schäffer, A. A. et al. VecScreen_plus_taxonomy: imposing a tax(onomy) increase on vector contamination screening. Bioinformatics 34, 755–759 (2017).

[46] De Nooy, W., Mrvar, A. & Batagelj, V. Exploratory Social Network Analysis With Pajek 3rd edn. (Cambridge University Press, 2018)

[47] Khan, A. W. et al. Super-Pangenome by Integrating the Wild Side of a Species for Accelerated Crop Improvement. Trends Plant Sci. 25, 148–158 (2020).

[48] Smith-Unna, R., Boursnell, C., Patro, R., Hibberd, J. M. & Kelly, S. TransRate: Reference-free quality assessment of de novo transcriptome assemblies. Genome Res. 26, 1134–1144 (2016).

[49] Aubry, S., Kelly, S., Kümpers, B. M. C., Smith-Unna, R. D. & Hibberd, J. M. Deep evolutionary comparison of gene expression identifies parallel recruitment of trans-factors in two independent origins of C4 photosynthesis. PLoS Genet. 10, e1004365 (2014).

[50] Simão, F. A., Waterhouse, R. M., Ioannidis, P., Kriventseva, E. V. & Zdobnov, E. M. BUSCO: Assessing genome assembly and annotation completeness with single-copy orthologs. Bioinformatics 31, 3210–3212 (2015).

[51] Waterhouse, R. M. et al. BUSCO Applications from quality assessments to gene prediction and phylogenomics. Mol. Biol. Evol. 35, 543–548 (2017).

[52] Nakamura, T., Yamada, K. D., Tomii, K. & Katoh, K. Parallelization of MAFFT for large-scale multiple sequence alignments. Bioinformatics 34, 2490–2492 (2018).

[53] Brown, N. P., Leroy, C. & Sander, C. MView: A web-compatible database search or multiple alignment viewer. Bioinformatics 14, 380–381 (1998).

[54] Kim, D., Song, L., Breitwieser, F. P. & Salzberg, S. L. Centrifuge: rapid and sensitive classification of metagenomic sequences. Genome Res. 26, 1721–1729 (2016)

[55] Breitwieser, F. P. & Salzberg, S. L. Pavian: interactive analysis of metagenomics data for microbiome studies and pathogen identification. Bioinformatics 36, 1303–1304 (2019)

[56] Kearse, M. et al. Geneious Basic: An integrated and extendable desktop software platform for the organization and analysis of sequence data. Bioinformatics 28, 1647–1649 (2012)

[57] NCBI Sequence Read Archive (SRA) https://identifiers.org/ncbi/insdc.sra:SRP229087 (2019).

[58] Luge, T., Fischer, C. & Sauer, S. Efficient application of de novo RNA assemblers for proteomics informed by transcriptomics. J. Proteome Res. 15, 3938–3943 (2016).

[59] Sievers, F. & Higgins, D. G. Clustal Omega for making accurate alignments of many protein sequences. Protein Sci. 27, 135–145 (2018).

